# Elucidating the functional profiles of cultivable bacterial communities in the water and sediments during the southwest monsoon season, along the west coast of India

**DOI:** 10.1101/2023.07.13.548871

**Authors:** Ashutosh S Parab, Mayukhmita Ghose, Cathrine S Manohar

## Abstract

Understanding the functional profiles of cultivable bacterial communities is crucial for comprehending their ecological significance in marine environments. This study investigates the functional profiles of cultivable bacterial communities in the water and sediments during the productive, southwest monsoon season, along the west coast of India. The southwest monsoon plays a vital role in shaping the hydrological and biogeochemical characteristics of this region, making it an ideal period to study bacterial communities and their functions. Our study utilizes a cultivation techniques to study taxonomy as well as functional assays to elucidate the diverse organic substrates utilization capabilities of bacterial communities. Cultivable bacteria were isolated from discrete water depths and sediment samples from the coastal and off-shore region. Subsequently the carbohydrase, lipase and protease assays were performed to assess its functional potential. The results of this study reveal a rich diversity of cultivable bacterial communities including, representative from Proteobacteria, Firmicutes and Actinobacterial phylum with diverse functional profiles. The functional analyses provide insights into the metabolic capabilities of the bacteria, including organic substrates degradation, processes. The bacterial taxonomic diversity and enzymes activities were significantly different (p < 0.001) among the water and sediment bacterial morphotypes. In conclusion, this research sheds light on the functional profiles of cultivable bacterial communities in the water and sediments along the west coast of India during the productive southwest monsoon season. The comprehensive analysis of their functional capabilities provides insights into their ecological roles and potential significance in organic matter recycling. These findings contribute to our understanding of microbial diversity and ecosystem functioning in coastal environments and lay the groundwork for further research on harnessing the significance of these bacteria in biogeochemical cycling.

## Introduction

The coastal oceans are extremely active, upwelling regions accounting for more than 10% of worldwide fresh output (Walsh 1991, Chavez & Toggweiler 1995, Wollast 1998, Gattuso et al. 1998). It is estimated that net phytoplankton production in the oceanic is around 50 Pg C per year, of which 10 Pg is exported to the ocean interior and the remaining 40 Pg is respired in the euphotic zone. Out of 10 Pg C, 8 Pg are known to be respired in the deep ocean, remaining 2 Pg ultimately reach the seafloor as organic carbon decomposition occurs while particles settle through the interior of the ocean (Middelburg, 2019). The organic matter degradation has a vast range of time scale and about 90% of the organic carbon is recycled within a few hours to days and only 0.2 Pg C yr-1 is eventually buried with a turnover of a millennia. Heterotrophic bacteria are the most prevalent and essential biological elements in the cycling of organic matter (OM) through the incorporation of carbon into their biomass and respiration (Hunter-Cevera et al. 2005). The distribution and production of bacterioplankton have attracted a lot of attention because they can serve as a vital link between dissolved OM and organisms at higher trophic levels (Azam et al., 1993). In marine ecosystems, heterotrophic bacteria are known to significantly contribute to microbial food webs and biogeochemical cycles by consuming up to half of the primary production (PP) through dissolved OM (Ducklow and Carlson, 1992; Azam et al., 1993; Lønborg et al. 2019). Bacteria primarily receive the dissolved organic substrates needed for their growth and biomass production from healthy growing phytoplankton, dead and decaying cells and through sloppy feeding by zooplankton (Fernandes and Bogati 2022). Bacterial extracellular enzymes initiate the microbial remineralization of phytoplankton-derived OM throughout the water column and sediment. Heterotrophic bacteria process a significant portion of marine primary productivity through extracellular enzyme activities. These enzymes catalyze the breakdown of high molecular weight OM to smaller substrates (about 600 Da) which can be effectively utilised for subsequent processing (Weiss et al. 1991). A portion of the carbon is integrated into biomass, another portion is converted to CO_2_, and the remainder is often expelled as refractory OM (Jiao et al. 2011). Thus, extracellular enzymes vary in their function in seawater and sediments, and the factors controlling their production, distribution, and activities are time-dependent and essential for carbon cycling in marine systems (Arnosti et al. 2011).

There is significant variation in the capacity of different bacterial groups to utilize OM. Also, these bacterial communities undergo seasonal fluctuations, thereby making the bacterial diversity and their associated enzyme activity studies even more crucial to get an enhanced understanding of their functional role in the ecosystem. The expression of these functional metabolic genes is unique at the species level and hence variation in bacterial community composition significantly alters their activity potential (Mahmoudi et al., 2020; Parab et al. 2022). Earlier studies have reported Proteobacteria, Firmicutes, and Actinomycetes as important bacterial groups from the marine environment, having diverse extracellular hydrolytic enzyme synthesizing potential and capacity to metabolize different substrates (Carbohydrates, proteins, lipids) based on the OM composition (Divya et al. 2010; Zimmerman et al. 2013; Teeling et al. 2016; Mahmoudi et al. 2020; Rocke et al. 2020). The availability of organic matter plays a crucial role in shaping the diversity and distribution of bacterial communities in both the water column and sediment. The presence of organic matter serves as a vital nutrient source for bacteria, influencing their growth and metabolic activity. Variations in organic matter availability, such as fluctuations in its quantity and quality, can lead to spatial variations in bacterial communities between the water column and sediment, as different niches provide distinct ecological conditions for bacterial colonization and function. In addition to organic matter availability, seasonal variations, including monsoons, upwelling, and other factors, have a profound impact on the physicochemical properties of a marine environment. These fluctuations subsequently trigger significant changes in bacterial diversity and functional enzyme activities. By altering factors like salinity, temperature, and nutrient availability, these seasonal variations shape the composition and behavior of bacterial communities.

The Arabian Sea is strongly influenced by physical forces such as winds, and heavy rainfall resulting in a significant increase in primary productivity during the southwest monsoon (Jyothibabu et al. 2010; Anjusha et al. 2018; Karnan et al. 2020). With significant upwelling and anthropogenic influence, the coastal regions surrounding the Arabian Sea are renowned for being extremely productive habitats and wide biodiversity in the World Ocean ecosystem (Gauns et al. 2005; Dalabehara and Sarma 2021; Parab et al. 2022). The hydrography of the Arabian Sea displays a distinctive characteristic primarily shaped by the seasonal shift of monsoonal winds and the consequent initiation of coastal upwelling, which typically starts in early June and continues till September (Gerson et al. 2014). The annual southwest monsoon triggers intense coastal upwelling in the Arabian Sea region i.e., the upward movement of nutrient-rich waters from sub-surface layers of the ocean. This phenomenon stimulates phytoplankton productivity and ultimately results in the production of a large amount of OM in this region (Ram et al. 2003; Jacob et al., 2009; Santhikrishnan et al. 2021). The nutrient-rich characteristics of upwelled water promote enhanced primary production, leading to increased bacterial abundance (Ramaiah et al. 2009; Lamont et al. 2014; Rocke et al. 2020). Apart from these physical forces, during the southwest monsoon, the Arabian Sea receives a significant influx of organic matter from anthropogenic sources, including rainfall and riverine runoff. The ultimate fate of this organic matter largely hinges upon the presence and metabolic activity of microorganisms within the marine system (Vipindas et al. 2020).

In recent years, there has been significant emphasis on exploring bacterial diversity through culture-independent approaches, utilizing advanced metagenomics techniques and Next Generation Sequencing (Alvarez-Yela et al. 2019; Vipindas et al. 2020; Sripan et al. 2021). However, despite these advancements, there is still a lack of knowledge concerning the metabolic activities and ecological significance of cultivable bacterial communities within this dynamic ecosystem. Culture-based diversity studies are important to gain deeper insights into the environmental realm of these bacteria, understand their functional metabolic processes in their natural habitat, and facilitate the detailed characterization of novel species (Mulla et al. 2018). In our previous research, we explored the diversity of cultivable and uncultivable microorganisms in this region during the monsoon season. Our findings indicated a greater bacterial diversity in the water column along the productive West coast of India (Parab et al. 2022 and Parab et al. 2023, Pre-print BioRivx). Building upon these findings, our current study focuses on comparing the bacterial diversity in both water and sediment during the productive monsoon season. Additionally, we aim to investigate the potential implications of this diverse biota on the degradation of organic matter. Also, a comprehensive investigation into the bacterial activity of the communities that thrive during the monsoon season during the upwelling is still lacking in this region. The present study aimed at understanding the effect of the southwest monsoonal on the cultivable bacterial diversity and their functional activities along the west coast of India. Also, this research brings in a comparative analysis of the bacterial activity potential of the bacterial morphotypes isolated from the water columns and sediment, thereby unfolding the ecological role and functioning of diverse bacteria in this dynamic productive zone during monsoon season.

## Methodology

### Study region and sampling

Water and sediment samples were collected along west coast of India (WCI), on the scientific cruise SSK-131 onboard RV Sindhu Sankalp during the monsoon season in September 2019. The sampling was conducted at three locations along WCI: off Goa, Mangalore, and Trivandrum (Fig.1), with a coastal and offshore station selected from each location. At each location, a coastal station located in the shelf region was positioned at a water depth of approximately 30 m, while an offshore station was positioned at a water depth of about 600 m. The coastal and offshore sampling stations situated off the coast of Goa were labeled as G30 and G600 respectively. The stations off Mangalore were titled as M30 and M600, and the Trivandrum station as T30 and T600 (Fig.1).

**Figure 1.**
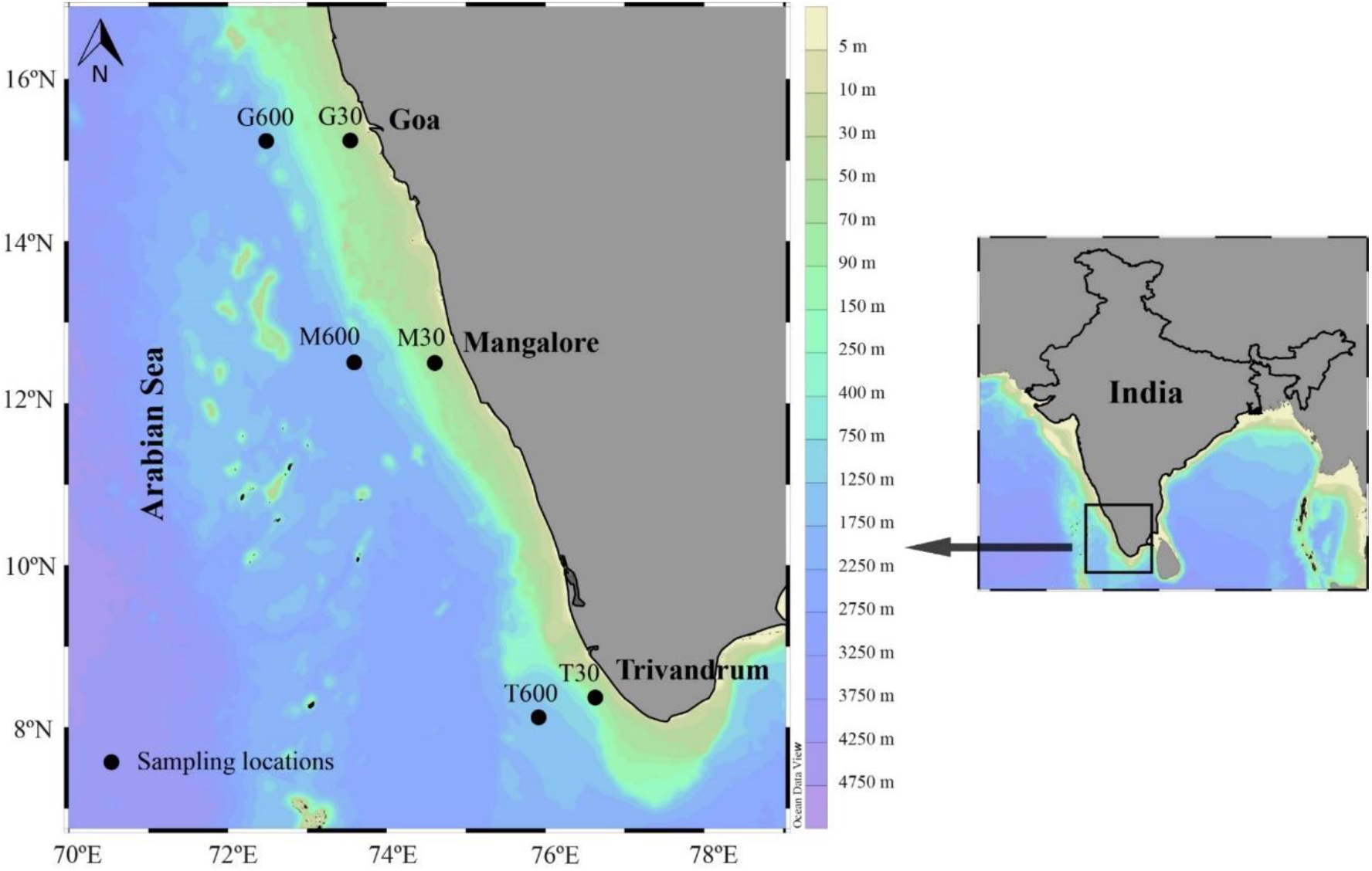
Study area and sampling stations along the west coast of India at Goa, Mangalore, and Trivandrum.

The underwater profiler (model: SBE-911 plus, Sea-Bird, USA) for conductivity-temperature-depth (CTD) was employed at all six stations to determine the physicochemical characteristics of the water column. Parameters including temperature, salinity, dissolved oxygen (DO), and fluorescence, which is indicative of the chlorophyll concentration were determined. At the coastal stations (G30, M30, and T30), water samples were obtained from the surface, chlorophyll maxima depth (Chl-M), and bottom waters. At the offshore stations, samples were collected from the surface, Chl-M, as well as depths of 100, 200, and 400 m, and from the bottom depth of 600 m. The chlorophyll maxima depth (Chl-M) varied depending on the station and was determined by analyzing the data acquired from the fluorescence sensor during the descent of the CTD sampling profiler. Subsequently, water samples were collected during the upward ascent using Niskin samplers (pre-cleaned before each sampling) attached to the underwater CTD unit (Sea-Bird, USA). At each of the locations, sediment samples were collected using a Peterson grab sampler. Subsequently, water and sediment samples were sub-sampled under aseptic conditions for further microbiological analysis.

### Assessment of bacterial abundance, viable counts and isolation of bacterial morphotypes

Bacterial abundance (BA) was assessed by adopting protocols from Porter and Feig (1980). For this, 20 mL subsamples of water collected from discreates depths were instantly fixed with 2% (final concentration) buffered formaldehyde after sampling and stored in the dark at ambient temperature for furthur analysis. The preserved samples were divided into multiple sets of 5 ml aliquots, with each set having three replicates. These aliquots were then incubated with 4’6’-diamidino-2-phenylindole (DAPI) at a concentration of 1 µg ml^-1^. Subsequently, the samples were passed through black polycarbonate membrane filters with a pore size of 0.22 µm (Millipore, U.S.A). The bacterial counts were determined using an epifluorescence microscope (Olympus BX-51, Japan) and reported as the number of cells per liter (cells L-1) of the water sample.

The total viable counts (TVC) of cultivable bacteria in both water and sediment samples were quantified using the spread plate method. To perform the enumeration, 100 µL aliquots of seawater samples and serially diluted 1 mg sediment samples were plated on Zobell Marine Agar (ZMA) plates in triplicate. The plates were then incubated under aseptic conditions at 28°C for a period of 24 to 48 hours. Suitable controls were included to prevent contamination and ensure accurate results. After the incubation period, the number of bacterial colonies was counted, and the results were expressed as colony-forming units (CFUs). The bacterial colonies were categorized into distinct morphotypes according to their colony appearance. For further analysis, a representative colony from each morphotype observed on the ZMA plates was streaked and isolated on new ZMA plate.

### Identification of bacterial morphotypes

The genomic DNA from each bacterial morphotype was extracted from pure cultures that were grown in Zobell Marine Broth (ZMB) for 24 hours at 28°C. The extraction process involved using a buffer with a high salt concentration to disrupt the bacterial cells and stabilize the DNA. This was followed by extraction with chloroform phenol and subsequent DNA precipitation using isopropanol. In order to identify the bacterial morphotypes, the extracted DNA from the pure cultures was amplified using PCR targeting the 16S rRNA gene. The amplification process involved the use of universal primers, namely 27F (5’AGAGTTTGATCMTGGCTCAG3’) and 1492R (5’TACGGYTACCTTGTTACGACTT-3’). (Parab et al. 2022). The amplified PCR products were subjected to sequencing using a cycle sequencing-ready reaction kit. The sequencing was performed on an Applied Biosystems (ABI) 3730 DNA stretch sequencer with the XL upgrade. The sequencing reaction utilized the ABI Prism BigDye terminator kit (version 3.1). Following the PCR amplification, the resulting DNA sequences were subjected to primer removal and quality check using Chromas software (version 2.6.6).

### Organic substrates utilization of bacterial morphotypes

The bacterial morphotypes isolated from the water and sediment samples were examined for their organic matter utilization potential using various carbohydrates, protein, and lipid substrates. The utilization of monosaccharides was estimated using liquid assays. For this, each bacterial morphotype was inoculated in a medium containing Bromocresol purple (pH indicator) and supplemented with monosaccharides such as glucose, arabinose, rhamnose, and galactose separately (Reiner 2012). The liquid media tubes were incubated for 48 h at 28°C. The ability of the bacterial morphotypes to utilize monosaccharides was evaluated by observing the color change of the liquid medium from purple to yellow, which indicates substrate utilization due to the subsequent pH shift in the medium. To measure the activity of extracellular enzymes involved in the hydrolysis of organic substrates such as polysaccharides (starch, carboxyl methylcellulose (CMC), agar, alginate, and xylan), basal salt agar plates were used. Each of the substrates mentioned was incorporated into the agar plates as a carbon source. The plates were then inoculated with the bacterial culture of interest, and the growth and activity were observed. The presence of clear zones or changes in the appearance of the substrate-containing agar indicated the hydrolysis of the respective polysaccharide, providing an estimate of the carbohydrase activity (Parab et al. 2022). Similarly, to assess the activity of extracellular enzymes involved in the degradation of organic substrates such as protein and lipid, basal salt agar plates were prepared. Specifically, Skim Milk (SM) was incorporated into the agar plates to evaluate protein degradation activity, while Tween 80 was added to assess lipid degradation activity (Fenice et al. 2007). The substrate plates were inoculated with bacterial morphotypes and then incubated for 48 h at 28°C. After the incubation period, the hydrolytic activity of amylase, agarase, and alginase enzymes were assessed by staining the agar plates with Gram’s iodine for a duration of 10 minutes. Cellulase and xylanase activities were determined by staining the agar plates with a 0.5% Congo red solution for a period of 15 minutes. After staining, the plates were destained by washing them with a 1N NaCl solution for 10 minutes. The carbohydrase activity was quantified by measuring the diameter of the zones of hydrolysis that had formed around the bacterial colonies after staining. To quantify the activity of proteases and lipases, the diameter of the zones of hydrolysis around bacterial colonies was measured. The hydrolytic enzyme activity (%) of the bacterial morphotypes within each phylum and genera was also estimated for all different substrates.

### Statistical analysis

Statistical analyses were conducted in the Microsoft Excel 2016 software. The Student’s t-test was employed to evaluate the difference between the distributions of hydrolytic activities exhibited by various bacterial morphotypes isolated from both water and sediment samples. To examine similarities among isolates’ activities based on bacterial taxonomy and isolation source, a dissimilarity matrix was constructed using the obtained results for each isolate. Non-metric multidimensional scaling (NMDS) was performed using the Vegan: community ecology (v2.6-4) and biodiversity: package for community ecology and suitability analysis (v2.15-2) in R Studio to explore these similarities. Furthermore, the significance of differences in the hydrolytic profiles of bacterial morphotypes, specifically about the activity levels for polysaccharides, proteins, and lipids, was investigated using the Permutational Multivariate Analysis of Variance (PERMANOVA) employing STAMP (statistical analysis of taxonomic and functional profiles) software. This analysis considered both the bacterial taxonomy and isolation source as factors to assess their contributions to the observed variations.

## Results

### Bacterial abundance and viable counts

The average BA in the water column of all coastal stations was estimated to be 54 ×10^9^ cells L^-1^, while BA in off-shore sites ranged from 41 to 57 × 10^9^ cells L^-1^. The average BA in the sediment samples collected in the coastal and off-shore station were in the range of 6 to 18 × 10^9^ cells L^-1^. Total viable counts estimated in the water column of coastal stations varied from 0.6 to 34×10^5^ CFU L^-1^ and it varied from 0.8 to 13 ×10^5^ CFU L^-1^ at the offshore stations. TVC levels in sediment samples from coastal stations reached as high as 27 to 57 × 10^7^ CFU L^-1^, whereas in offshore stations, average counts were estimated to be 22 to 36 × 10^7^ CFU L^-1^. TVC levels in sediment were higher than in the water column. There was no visible trend observed in the TVC counts (Table 1).

**Table 1.** Distribution of bacterial abundance (BA) and total viable counts (TVC) in the water column and sediment in coastal and offshore stations.

| Station | Bacterial abundance |  | Total viable counts |  |
| --- | --- | --- | --- | --- |
|  | Water | Sediment | Water | Sediment |
|  | (10<sup>9</sup> cells L<sup>-1</sup>) | (10<sup>9</sup> cells L<sup>-1</sup>) | (10<sup>5</sup> CFU L<sup>-1</sup>) | (10<sup>7</sup> CFU L<sup>-1</sup>) |
| **Coastal stations** |  |  |  |  |
| **G30** | 53 | 11 | 34 | 27 |
| **M30** | 56 | 13 | 10 | 42 |
| **T30** | 54 | 15 | 0.6 | 57 |
| **Offshore stations** |  |  |  |  |
| **G600** | 57 | 18 | 13 | 36 |
| **M600** | 41 | 6 | 15 | 23 |
| **T600** | 44 | 8 | 0.8 | 22 |

### Bacterial morphotypes identified from water and sediment

A total number of 92 and 59 morphotypes from the water and sediment samples respectively, were subjected to DNA isolation and sequencing. Diversity analysis was conducted by characterizing unique cultivable morphotypes using 16S rRNA sequencing. The bacterial sequences obtained were subjected to analysis in the NCBI database using the bioinformatics tool BLAST (Basic Local Alignment Search Tool) for confirmation of their taxonomic and phylogenetic resemblance. The sequences corresponding to the bacterial morphotypes derived from water and sediment samples have been deposited in the database with the following GenBank accession numbers: OP198814 – OP198905 for water samples and OP168205 – OP168263 for sediment samples. Morphotypes from the water and sediment samples belonged to the major phyla Proteobacteria, Firmicutes, and Actinobacteria. The relative abundance of Firmicutes phylum was highest (71 %), followed by Proteobacteria (26 %) and Actinobacteria contributing 1 % each in the water samples (Fig. 2). In sediment samples, the bacterial communities included Firmicutes, which was represented with the highest relative abundance of 71 % followed by Proteobacteria (22%), and Actinobacteria (7 %).

**Figure 2.**
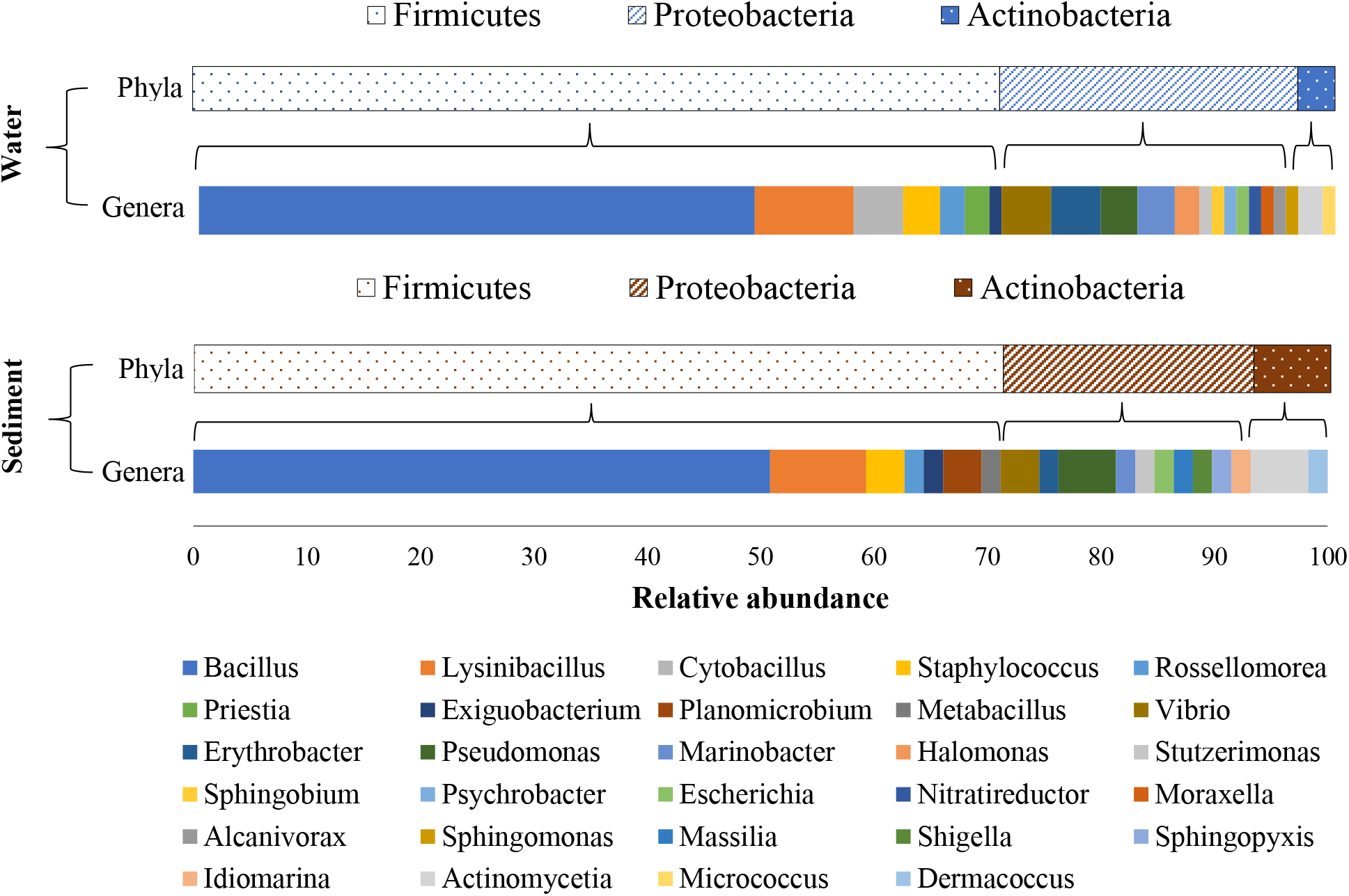
Taxonomic composition of cultivable bacterial morphotypes at phyla and genera levels from water and sediment samples along the coastal and offshore stations during the September 2019 sampling season.

The molecular characterization of the 151 morphotypes at the genus level obtained from both water and sediment samples. The morphotypes were categorized into 29 different genera, with 22 genera identified in the water samples and 19 in the sediment samples. Interestingly, 12 genera were found to be common to both sample types. Further analysis of the water samples revealed that the bacterial morphotypes could be classified into 13 genera belonging to the Proteobacteria phylum, specifically the alpha (4 genera) and gamma (9 genera) class. Additionally, the phylum Firmicutes was represented by 7 genera, while the Actinobacteria phylum was represented by only 2 genera.

Bacterial genera *Vibrio* and *Erythrobacter* belonging to Proteobacteria, *Bacillus,* and *Lysinibacillus* of phylum Firmicutes were dominant in the water samples (Fig. 2). Actinobacteria were represented by *Actinomycetia* and *Micrococcus* genera in water samples. In the sediment samples, Proteobacteria were represented by 10 genera, of which 2 were from Alpha, 1 and 7 were from Beta and Gammaproteobacteria classes respectively. Firmicutes were represented by 7 genera, and Actinobacteria were represented by 2 genera. *Pseudomonas* and *Vibrio* from Proteobacteria and Bacillus and Lysinibacillus from Firmicutes were dominant in sediment samples. Actinobacteria morphotypes were represented by 2 genera including *Dermacoccus* and *Actinomycetes* in the sediment samples (Fig. 2).

### Organic substrates utilization ability of bacterial morphotypes

All the representative 92 morphotypes from water samples and 59 morphotypes from sediment samples were subjected to characterization of hydrolytic enzyme activity. The results demonstrated that the bacterial morphotypes exhibited enzymatic activity for all three major substrate groups, including carbohydrases, proteases, and lipases. Interestingly, it was observed that the bacterial morphotypes showed a preference for monosaccharide glucose as the most utilized substrate among the different carbohydrates tested. This indicates the ability of these bacteria to efficiently metabolize glucose as a carbon source.

(Fig. 3). The bacterial morphotypes from sediment samples showed higher hydrolytic enzyme activity. Among total morphotypes, approximately 50 % showed good activity in all the substrates except rhamnose. Based on the zone of hydrolysis, the percentage of morphotypes from the water samples showed activity in alginate > agar > starch > CMC > xylan > SMA. Morphotypes isolated from sediment samples showed activity in agar > starch > alginate > CMC > xylan > Tween 80 (Fig. 3). Monosaccharide glucose, polysaccharides starch, agar, and alginate lyase activity were higher as compared to other substrates. The percent activity of bacteria isolated from sediment samples was comparatively higher than bacteria isolated from water samples (Fig. 3). The hydrolytic enzyme activity of the morphotypes belonging to each phylum and genera level was studied and the results show that bacterial isolates from Phylum Firmicutes followed by Proteobacteria from both the samples exhibited good hydrolytic activity and were able to utilize all the 11 substrates tested. Firmicutes isolated from water samples showed higher activity in alginate (85 %) followed by agar (82 %) starch (72 %) and CMC (71 %). (Fig. 4).

**Figure 3.**
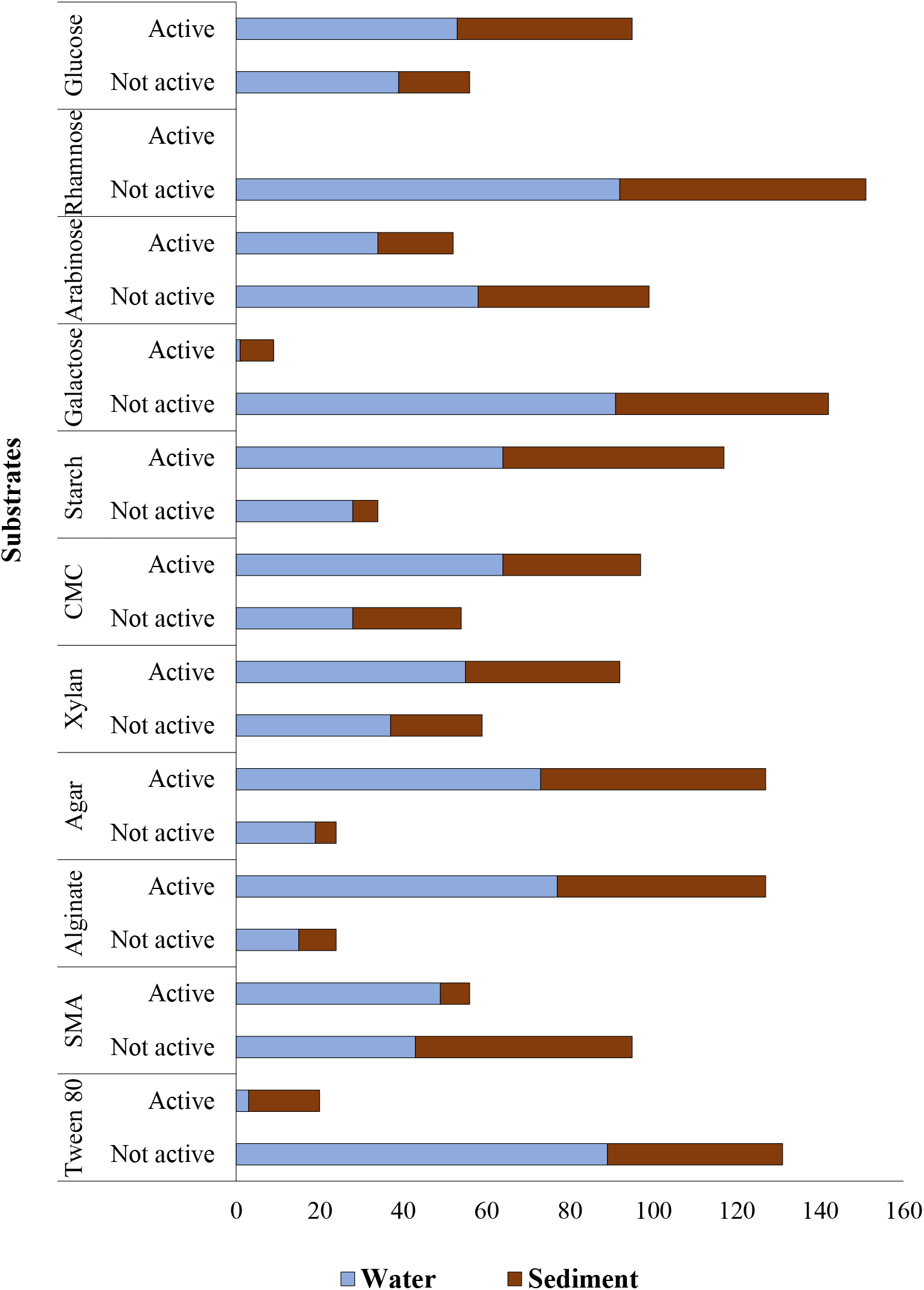
Hydrolytic activities of bacteria isolated from the water and sediment samples. Numbers of active and non-active bacteria in the water and sediment samples along the coastal and offshore stations.

**Figure 4.**
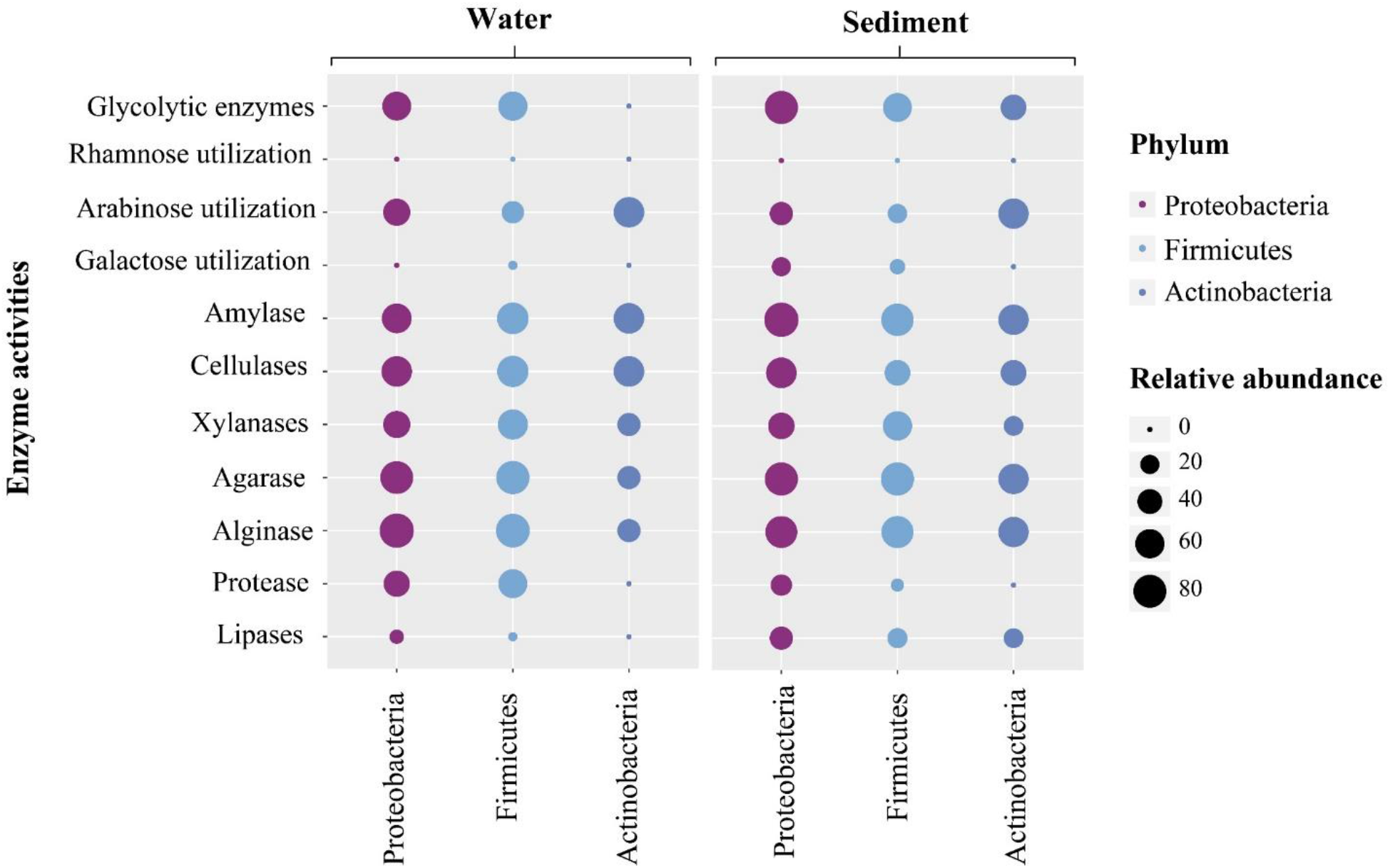
Bubble chart of the tested hydrolytic enzyme activity (%) of bacterial morphotypes of different phyla isolated water and sediment samples along the coastal and offshore stations along the west coast of India.

Within phylum Firmicutes, *Bacillus* spp. showed higher activity across all substrates compared to other genera. Bacterial genera *Exiguobacterium* spp. *Staphylococcus* spp. and *Cytobacillus* spp. also showed marginally higher utilization activities. Proteobacteria also showed good activity for alginate (88%), agar (79 %), alginate (67 %), and starch (63%). Actinobacteria showed good activity in Monosaccharides arabinose (67 %), polysaccharides starch (67 %), and CMC (67 %) (Fig. 4). Proteobacterial genera *Vibrio* spp., *Sphingomonas* spp., *Moraxella* spp., *Marinobacter* spp. and *Alcanivorax* spp. were among the dominant genera which showed higher utilization activities. Actinobacteria were represented by *Actinomycetia* sp. and *Micrococcus* sp. showed marginally lower hydrolytic activities compared to other taxa.

Bacterial morphotypes isolated from sediment samples showed marginally higher activity as compared to bacterial morphotypes isolated from water samples. Although the bacterial phyla from water and sediment samples were similar, the bacterial activities at the genera level were different within each phylum and community shift in genera level was also observed based on activity tested. All Proteobacterial morphotypes showed activity in starch followed by glucose (92 %), agar (92 %), and alginate (85 %). Among Proteobacteria, genera including *Pseudomonas* spp. *Vibrio* sp. *Erythrobacter* spp. and Escherichia spp. showed higher utilization activities in almost all substrates tested. Bacterial morphotypes belonging to Firmicutes showed good activity in agar (93 %), starch (88 %), and alginate (86 %). Bacteria genera including *Bacillus* spp., *Lysinibacillus* spp. and Staphylococcus spp. were among the dominant genera contributing to overall activity in the Firmicutes phylum (Fig. 4 & 5). Actinobacterial morphotypes isolated from sediment showed comparatively higher activity than water. Agar, alginate, and arabinose activity showed by Actinobacteria was comparatively higher than other substrates (Fig. 5). Bacterial genera including *Actinomycetia* spp. and *Dermacoccus* spp. were the only dominant taxa observed and contributing to the organic substrates utilization activities among the phylum Actinobacteria.

**Figure 5.**
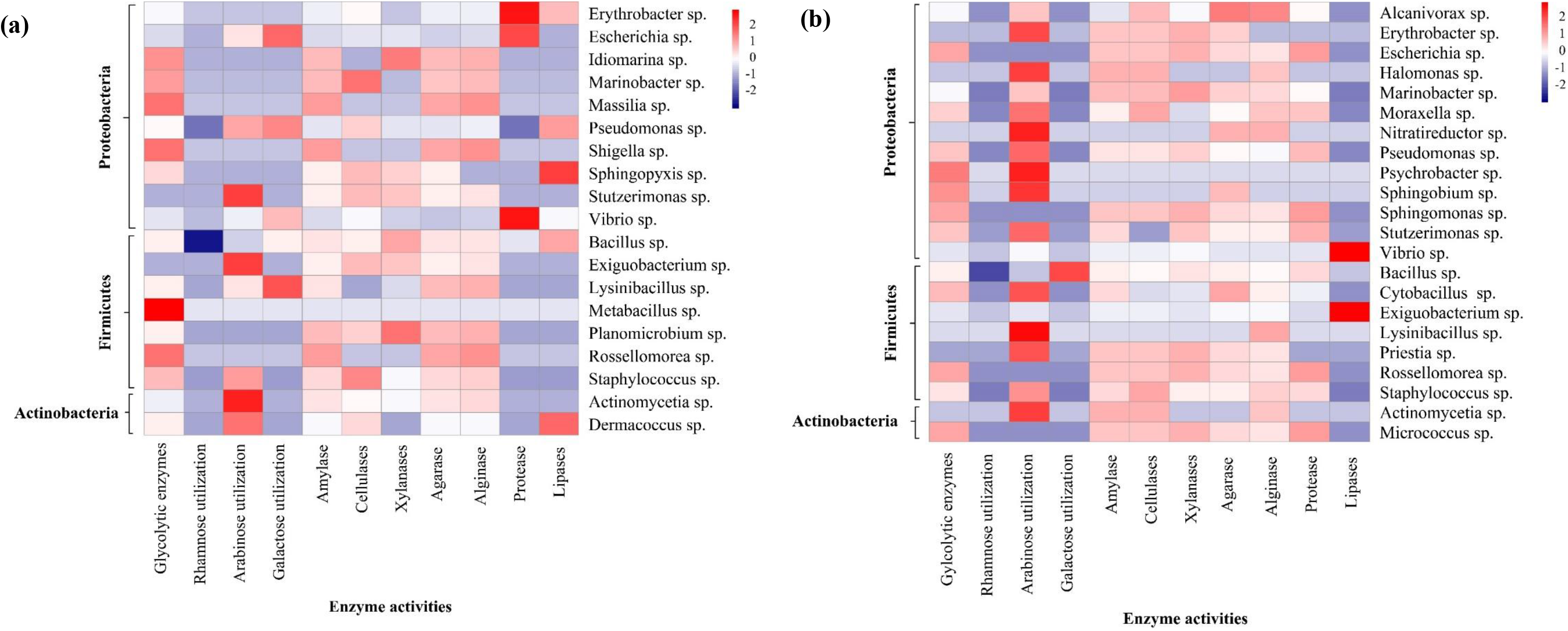
Heatmap of the tested hydrolytic enzyme activity (%) of bacterial morphotypes of different genera isolated from water (a) and sediment (b) samples along the coastal and offshore stations along the west coast of India.

Based on the activity level (hydrolysis zone in substrates medium), the top ten bacterial morphotypes from water and sediment samples showing higher activities in more than five substrates were also evaluated. Among this top five bacterial morphotypes isolated from water samples were phylogenetically affiliated to *Halomonas axialensis*, *Vibrio natriegens, Bacillus cereus, Halomonas axialensis, Bacillus thuringiensis and Lysinibacillus boronitolerans* were from the dominant bacteria with overall higher activity levels noted (Supp. Fig. 1 Fig. 6). In sediment samples, bacterial morphotypes affiliated to *Halomonas axialensis*, *Vibrio natriegens* and *Bacillus thuringiensis* showed the highest alginase and agarase activity with activity zone values around 30 ± 2 mm. *Lysinibacillus* sp. showed the highest activity (24 ± 2 mm) for xylanase, agarase and protease. Bacteria isolated from sediment including *Dermacoccus profundi, Erythrobacter citreus, Escherichia coli, Bacillus albus* and *Bacillus pumilus* were the most active in degrading the majority type of carbohydrates, proteins and lipid substrates. *Erythrobacter citreus, Bacillus albus*, and *Dermacoccus profundi* showed higher activity zone in Agar (30 mm) (Fig. 6)

**Figure 6.**
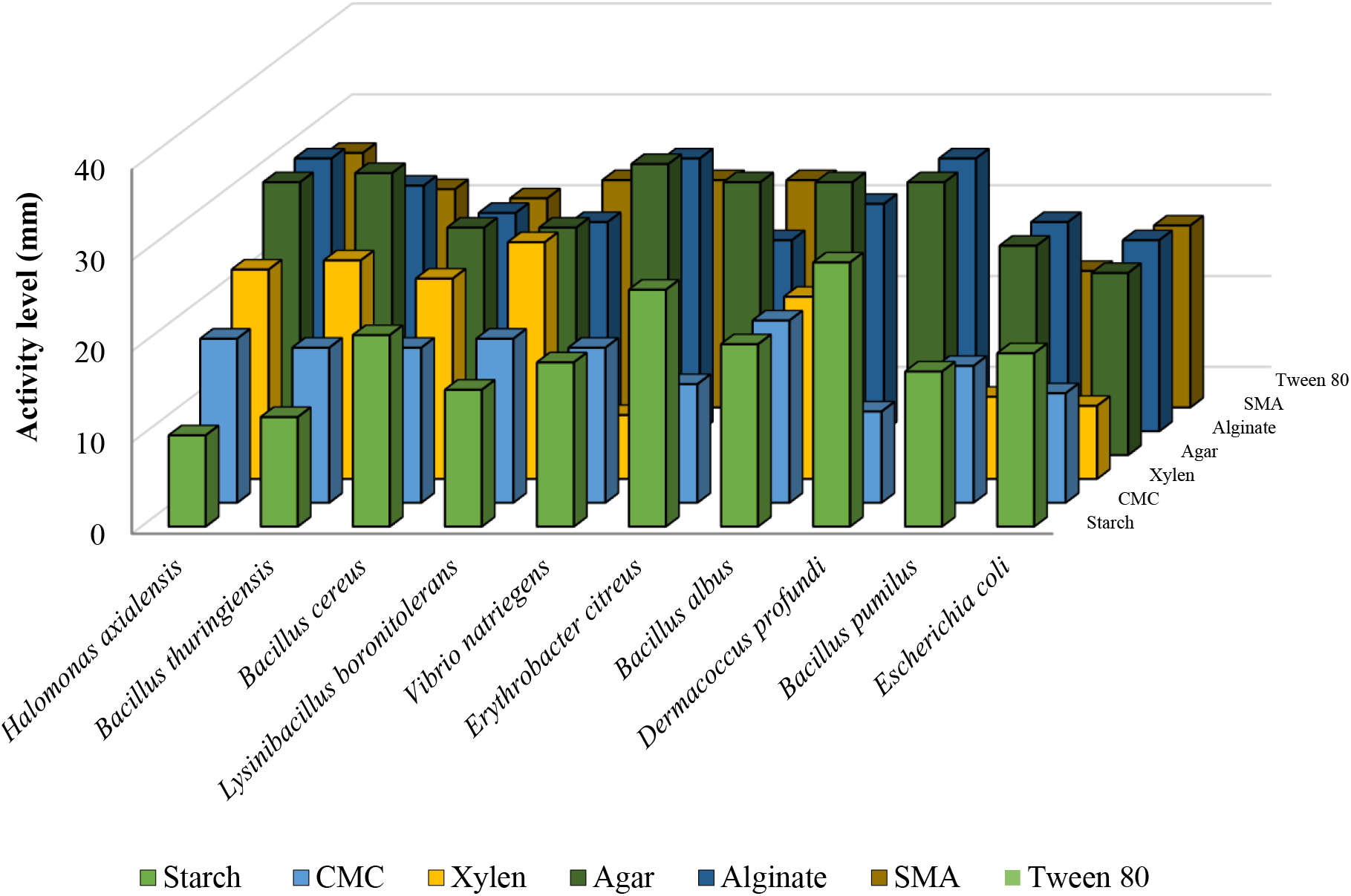
Bacterial morphotypes with overall higher activity levels. Activity levels were reported as the zone of hydrolysis (diameter in mm) around bacterial colonies isolated from the water and sediment with the highest activities.

### Comparative analysis of substrates utilization pattern

The Student’s t-test was employed to assess the difference between the distributions of hydrolytic activities (zone in mm) exhibited by various bacterial morphotypes isolated from both water and sediment samples. The difference between the activity of the level of bacterial morphotypes isolated from water and sediment was significantly different (Table 2). For amylase activity, the mean activity level in water is 10.9 mm, while in sediment, it is slightly lower at 9. The t-test indicates a statistically significant difference (p < 0.05), suggesting that the amylase activity differs between water and sediment environments. Protease activity also demonstrates comparable mean activity levels in water (17.1 mm) and sediment (17.7 mm), suggesting a significant difference between the two samples. Similarly, lipase activity displays a mean activity level of 9.7 in water with an SD of 3.5, while in sediment, it is slightly higher at 11.9 with an SD of 4.8. The t-test indicates a statistically significant difference (p < 0.05), implying that lipase activity differs between water and sediment (Table 2). However, the difference between cellulase, xylanase, agarase and alginase activities of bacterial morphotypes isolated from water and sediment were not significant. These results suggest significant variations in the activities of certain enzymes, such as amylase, protease, and lipase, between the water and sediment morphotypes, indicating potential differences in the degradation processes and microbial communities associated with each substrate.

**Table 2.**
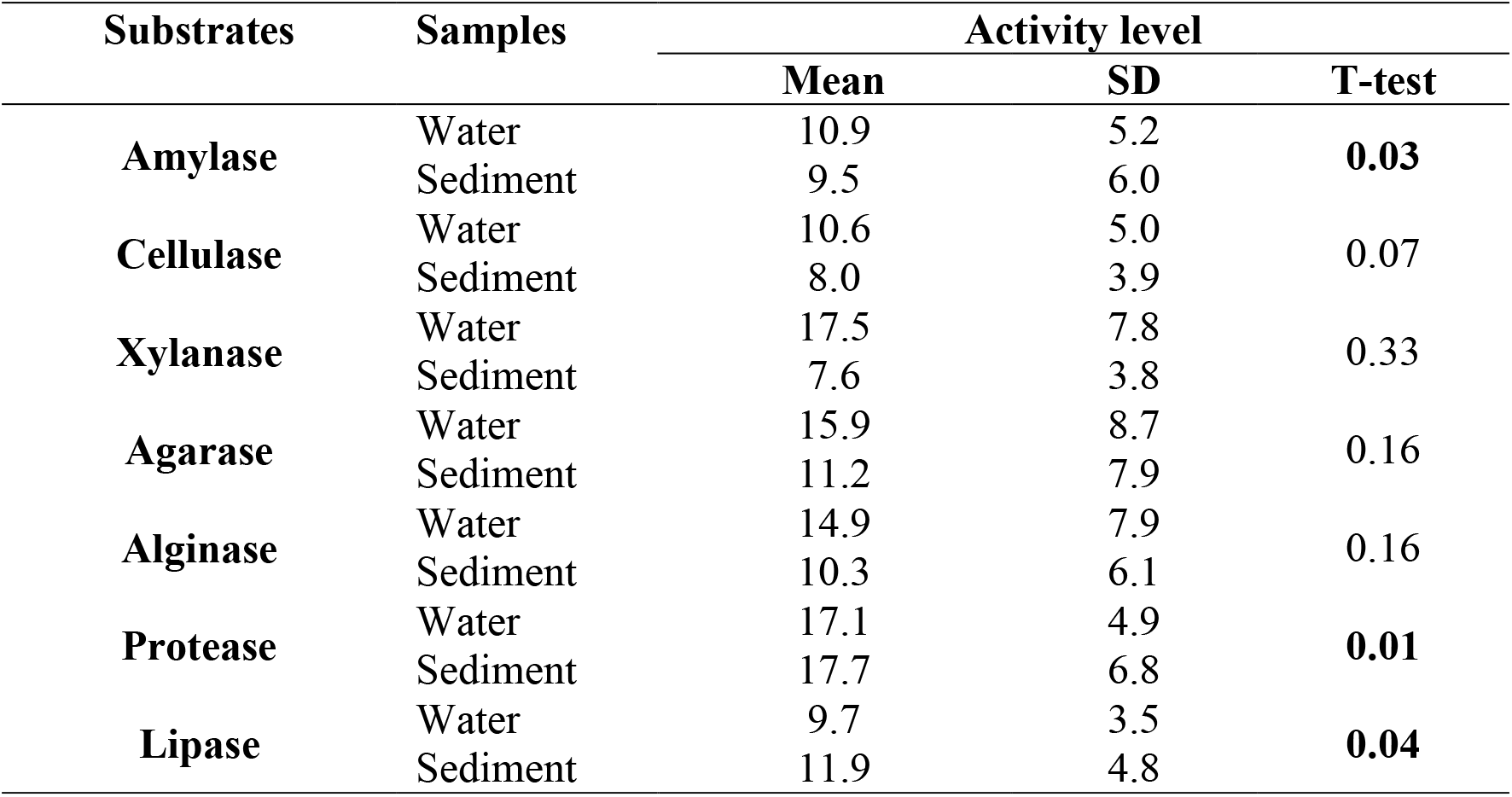
Significant distribution of hydrolytic activities in the water and sediment. In bold: p < 0.05

To examine similarities among isolates’ activities based on bacterial taxonomy at the phyla level and isolation source, Non-metric multidimensional scaling (NMDS) was performed (Supp Fig. 2). Furthermore, these significances of differences in the hydrolytic profiles of bacterial morphotypes, specifically concerning the activity levels for polysaccharides, protein, and lipid, was also investigated using the PERMANOVA (Table 3). This analysis considered both the bacterial taxonomy and isolation source as factors to assess their contributions to the observed variations. A total of 9999 permutations were performed to assess the significance of the observed differences. The results demonstrated a significant difference between groups when water and sediment morphotypes were compared together. At the phylum level, significant differences were observed (p = 0.0001) between groups, while at the genus level, a similar pattern was observed (p = 0.002). Furthermore, the analysis examined the microbial composition within water and sediment samples separately. For water samples, although no significant differences were found at the phylum level, a highly significant difference was observed at the genera level (p = 0.007), indicating some dissimilarity in phylum composition within water morphotypes at the genera level. Similarly, sediment morphotypes at the phyla level did not show any significant difference, however, at the genera level variations were marginally significant (p = 0.04) (Table 3). The PERMANOVA analysis revealed significant differences in microbial composition and enzyme activities among the water and sediment groups examined.

**Table 3.** PERMANOVA test performed on water, sediment and overall morphotypes at phylum and genera level based on activity. In bold: p < 0.05

|  | Groups | Permutation<br>N: | Total sum<br>of<br>squares: | Within-<br>group sum<br>of squares: | F: | p<br>(same): |
| --- | --- | --- | --- | --- | --- | --- |
| **Overall** | Phylum | 9999 | 21.3 | 19.7 | 10.9 | **0.0001** |
|  | Genus | 9999 | 21.3 | 14.0 | 1.7 | **0.0002** |
| **Water** | Phylum | 9999 | 13.2 | 12.6 | 2.1 | 0.07 |
|  | Genus | 9999 | 13.2 | 7.5 | 1.6 | **0.007** |
| **Sediment** | Phylum | 9999 | 6.5 | 6.0 | 2.0 | 0.05 |
|  | Genus | 9999 | 6.5 | 3.5 | 1.5 | **0.04** |

## Discussion

### Hydrography

Physicochemical characteristics determined from coastal and offshore sampling locations along the WCI showed spatial variability in this region. Sea surface temperature in coastal and offshore stations averaged 28°C with a 1°C variation. Salinity remained consistent at 33 ± 3 PSU in coastal stations but varied in offshore stations. Dissolved oxygen levels were 4 ± 0.5 mg L^-1^, dropping at mid-depth in offshore stations. Chlorophyll concentrations based on fluorescence sensor were highest at the surface waters and decreased with depth. Nitrate concentrations averaged 4.9 µM in coastal stations and 23.6 µM in offshore stations. Nitrite levels were generally low, except in one coastal station. Phosphate and silicate levels showed variations between coastal and offshore stations. The comprehensive analysis and interpretation of the physicochemical conditions of the water column examined in this study can be found in the pre-print article (Parab et al. 2023, available on BioRix). The analysis of physicochemical parameters indicates that the water column condition exhibited characteristics of upwelling phenomenon (Dalabehara et al. 2021). This was observed through a shallower mixed layer depth and an increase in subsurface chlorophyll and nutrient concentrations. This processes are fundamental component of primary production in this region (Habeebrehman et al. 2008). The increased phytoplankton biomass resulting from upwelling fuels the subsequent stages of the marine food web. The accumulation of organic matter derived from the primary production, including dead phytoplankton and other organic detritus in the water column and subsequently into the sediment creates a favorable environment for bacterial proliferation. The bacterial proliferation stimulated by the availability of organic matter lead to an increase in microbial activity and biomass, contributing to the overall nutrient cycling in this upwelling region. These processes create a dynamic and interconnected ecosystem where the generation of organic matter through primary production supports the growth and abundance of bacteria, forming an essential link in the marine food web and nutrient recycling.

### Variation in the diversity of bacterial morphotypes isolated from water and sediment

Through the application of 16S rRNA sequencing, the evaluation of bacterial diversity in both water and sediment within the productive upwelling zone of the Arabian Sea revealed the presence of bacterial morphotypes belonging to three prominent phyla: Firmicutes Proteobacteria and Actinobacteria. Previous studies have consistently identified the phyla Proteobacteria, Firmicutes, and Actinobacteria as the predominant bacterial groups in this region. This finding further supports the notion that these phyla play a crucial role in shaping the bacterial composition and dynamics within the productive upwelling zone of the Arabian Sea (Amberkar et al. 2021; Parab et al. 2022). Firmicutes contributed maximum during the monsoon season showing an abundance of 71% each in both water and sediment. However, although the relative abundance of Firmicutes was significantly high in monsoon, the diversity of bacterial morphotypes at the genus level was significantly low, with *Bacillus* spp. as the major genera present. This can probably be due to the increased phytoplankton production during the monsoon season and their associated synthesis of anti-bacterial compounds, which largely restricted the growth of another major genus of Firmicutes such as *Staphylococcus*, which are prevalent in other seasons (Shannon and Abu-Ghannam 2016; Mühlenbruch et al. 2018; Isaacs et al. 2021). Such conditions also favored the growth of bacteria of the genus *Micrococcus* sp., and *Actinomycetia* sp. of phylum Actinobacteria in this season.

Proteobacteria, which is a prominent phylum, is widely found in different marine ecosystems including coastal regions (Khandeparker et al. 2017), upwelling zones (Li et al. 2014; Vijayan et al. 2021), and oxygen minimum zones (OMZ) (Bandekar et al, 2018; Mulla et al. 2018; GuerreroFeijóo et al. 2017; Vijayan et al. 2021). During monsoon, Proteobacteria contributed to maximum diversity for both water and sediment samples. However, there were significant differences in their representative morphotypes. For water samples, only morphotypes belonging to Alpha and Gammaproteobacteria were found, while beta Proteobacteria were not reported from any of the study regions. The complete absence of beta Proteobacteria in the water column during monsoon was in good accordance with the earlier study (Parab et al. 2022) in the Arabian Sea region. However, Proteobacteria showed all three alpha, beta, and gamma morphotypes in the sediment sample, with beta Proteobacteria contributing the least. Among Proteobacteria, *Vibrio* spp. was one of the dominant Proteobacterial genera found in both water and sediment of this region. Despite showing wide diversity, the abundance of phylum Proteobacteria was significantly less in the monsoon season, as compared to the other seasons studied from this region. This results were in good accordance previous studies (Vijayan et al. 2021, Parab et al. 2022).

A marginal increase in Actinobacteria abundance was observed in sediment samples compared to water samples. Actinobacteria, a diverse phylum of bacteria, play a vital role in marine ecosystems. Sediments provide a stable and nutrient-rich environment, favoring Actinobacteria growth (Behera et al. 2017). Actinobacteria play a crucial role in the decomposition of complex organic biopolymers, such as lignocellulose and chitin. This ability makes them significant contributors to the cycling of organic matter in sedimentary environments (Lv et al., 2014). Seasonal variations and increased organic matter and nutrient inputs during the monsoon may contribute to this observed pattern. The water and sediment samples in this study were found to be predominantly represented by Actinomycetia at the genus level. The prevalence of Actinomycetia in both water and sediment samples suggests their significant contribution to the microbial community composition in this particular environment. Actinomycetia genera are known for their diverse metabolic capabilities and ecological roles, including the production of bioactive compounds and participation in nutrient cycling processes (Jagannathan et al. 2021).

### Variation in substrates utilization pattern of bacterial morphotypes isolated from water and sediment

Bacteria are the key players in organic matter (OM) recycling in the ocean. They utilize the major natural polymers of OM, such as carbohydrates (especially polysaccharides, and a few monosaccharides), proteins, and lipids by producing different hydrolytic enzymes (Parab et al. 2022). There are few investigations of bacterial hydrolytic enzyme activities conducted in different coastal ecosystems along west coast of India (Baby Divya 2010; Khandeparker et al. 2011; Singh and Ramaiah 2011; Parab et al 2022). The studies have shown that bacterial enzyme activity varies significantly with seasons and is specific to different ecological zone, owing to varying OM composition, making such studies significantly important (Basu et al. 2013; Khodse and Bhosle 2013; Mahmoudi et al. 2020). Hence, to the best of our knowledge, the present study presents a firsthand comparative analysis of the hydrolytic enzyme activities between cultivable bacteria derived from water and sediment samples collected from the coastal upwelling region of the Arabian Sea. This research provides valuable insights into the significance of bacterial diversity and its functional role in the ecosystem, especially during the southwest monsoon season. A substrate-based activity study was performed to determine the enzyme hydrolytic potential of cultivable bacteria in both water and sediment from the productive upwelling ecological zone during monsoon. Bacterial isolates of the phylum Firmicutes showcased the highest enzyme activity and metabolic diversity, with the capacity to utilize all eleven substrates under investigation. Although isolates from the phylum Proteobacteria showed similar metabolic diversity to Firmicutes, enzyme activity was lower than the Firmicutes as their abundance was significantly low in this season. For Actinobacteria, metabolic diversity was significantly less, in both sediment and water samples, showing enzymatic activity against five out of eleven substrates tested. A similar pattern of enzyme activity was observed for Actinobacteria from the OMZ sediments of the Arabian Sea (Divya et al. 2010).

Bacterial hydrolytic enzyme activity was significantly found to be higher in sediment samples in this study region as compared to water. Marine sediments serve as a crucial repository for nutrients derived from both natural processes occurring in the water column and anthropogenic activities. The abundance of nutrients makes sediments an ideal habitat for a wide range of bacterial species (Fierer and Lennon, 2011; Vipindas et al. 2020). The microbial activity in the sediment plays a vital role in replenishing nutrients to the water column, which is essential for regulating the ecological functioning of several biogeochemical cycles in the ocean and maintaining the overall biological productivity of the Arabian Sea. The Arabian Sea is a highly productive marine ecosystem that is constantly changing due to various environmental factors. It experiences seasonal influences from physical forces like upwelling, winter cooling, and monsoonal winds. The southwest monsoon brings in a significant amount of anthropogenic waste from riverine runoff and terrestrial sources, which largely increases the organic matter content of the ecosystem (Vipindas et al. 2020). Additionally, the monsoon season promotes the upwelling of nutrient-rich subsurface waters, creating favorable conditions for elevated biological productivity and significant accumulation of organic matter in the bottom sediment (Kurian et al. 2020). A considerable portion of organic matter, whether generated within the water column or brought from other sources, eventually settles into the sediments at the bottom of the seafloor, creating a rich reservoir of nutrients (Vipindas et al. 2020). This can probably be the reason for higher bacterial diversity leading to higher enzyme expression and activity in sediments favoring the utilization and recycling of nutrients throughout the water column. Of the 11 substrates tested in this study, monosaccharide glucose was found to be the most utilized substrate for both water and sediment. The study by Parab et al. 2022 also found glucose being the most hydrolyzed monosaccharide in the Arabian Sea across the seasons. Also, Lipase activity was mainly observed in sediment samples. Earlier studies have reported that deep-sea microorganisms utilize lipids as their major nutritional source, which are mainly associated with sediments. Thus, sediment-associated bacteria express a significant amount of lipase enzymes, resulting in higher lipase enzyme activity in sediments (Schulte et al. 2000; Baby Divya et al. 2010). Based on the activity zone, top ten morphotypes from water and sediment showing highest activity, were also reported from various marine region for their higher bioactive potential to utilize various carbohydrates, proteins and well as lipid containing substrates (Supp table 1).

The conventional methods of cultural isolation and sequence-based identification of microorganisms from natural samples are still important for understanding their detailed functional role and biodiversity in the environment. However, one of the major limitations of cultural techniques is that a large proportion of cultures from marine environments are viable but non-cultivable, and do not express in culture media and therefore cannot be studied through conventional methods of cultivation. Despite the disadvantages, culture-based studies can provide a better understanding of the natural habitat of the bacteria (Amberkar et al. 2021; Parab et al. 2022). Also, this traditional method can specifically pose an advantage to bacteria being present in very low concentrations in the environment, which often gets undetected in an advanced metagenomic study of culture-independent isolation of bacteria from environmental samples (Sanz-Sáez et al. 2020).

## Supplementary Figures and tables

**Supplementary Figure 1.**
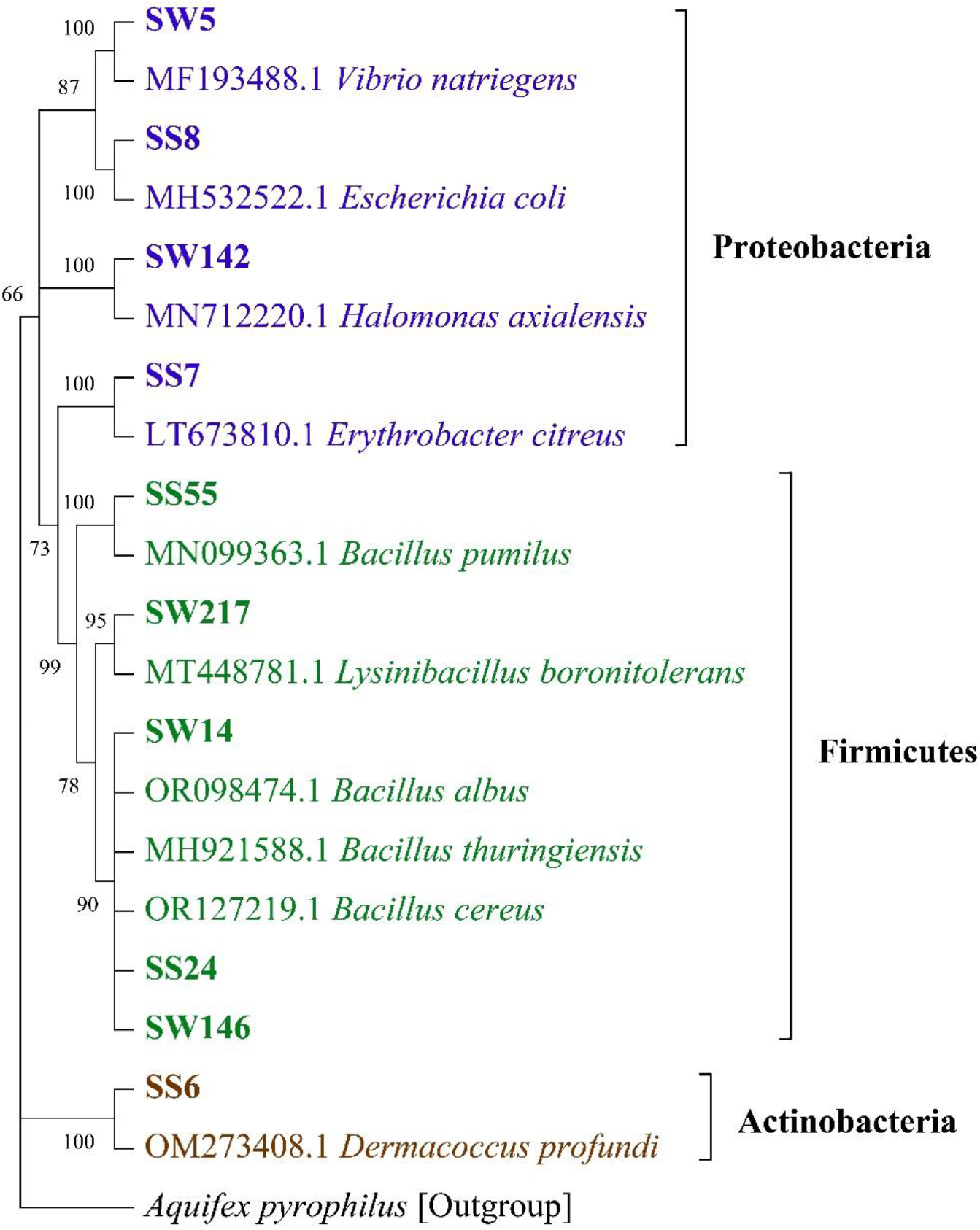
Maximum-likelihood-based phylogenetic tree of 16 rRNA gene sequences of top ten bacterial morphotypes from water and sediment with higher substrate utilization activity.

**Supplementary Figure 2.**
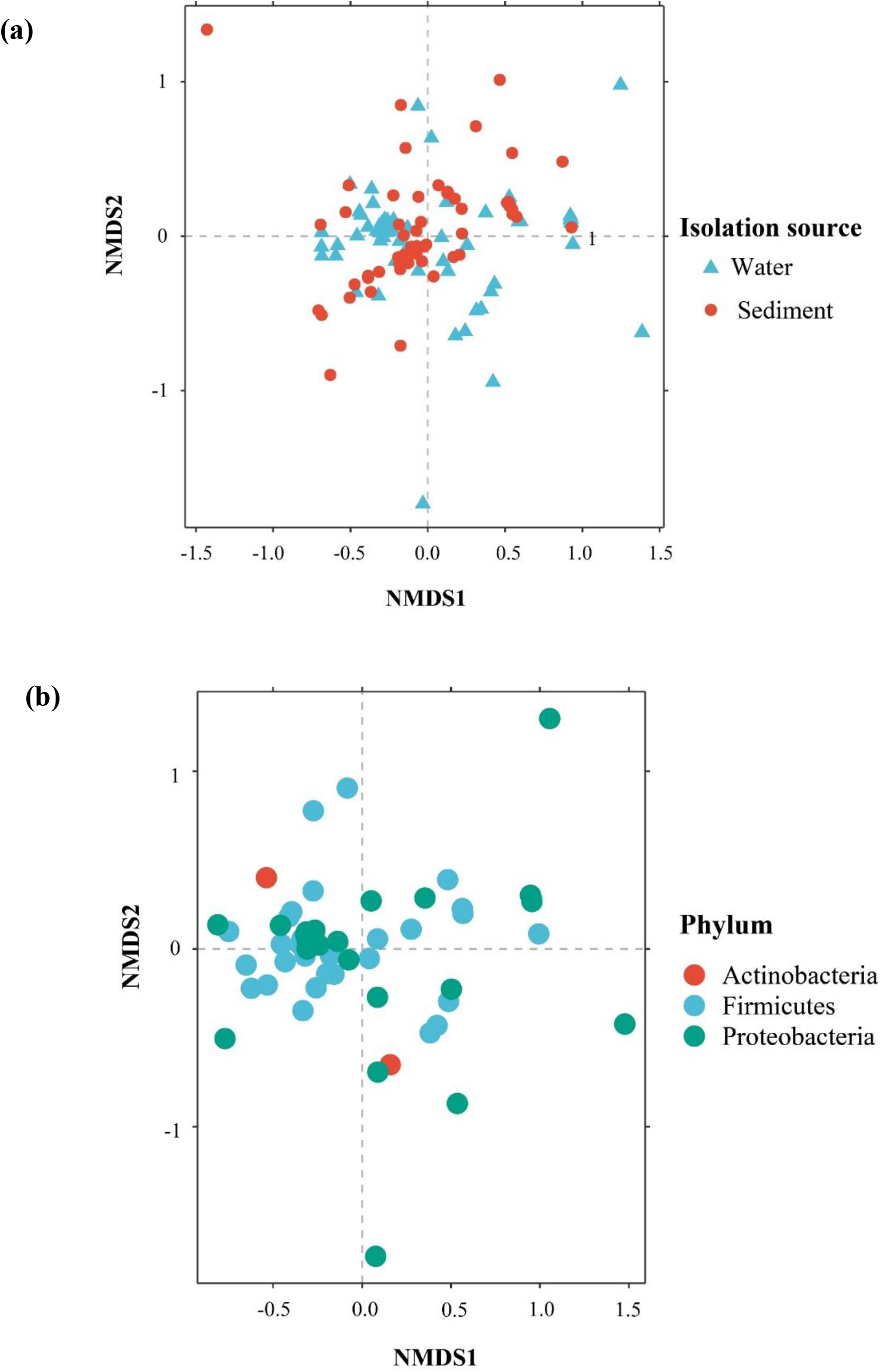
Non-metric MDS distribution of the morphotypes based on the tested polysaccharides, protease, and lipase activities. Morphotypes are distinguished based on the (a) isolation source and (b) bacterial phyla level.

**Supplementary Table 1.**
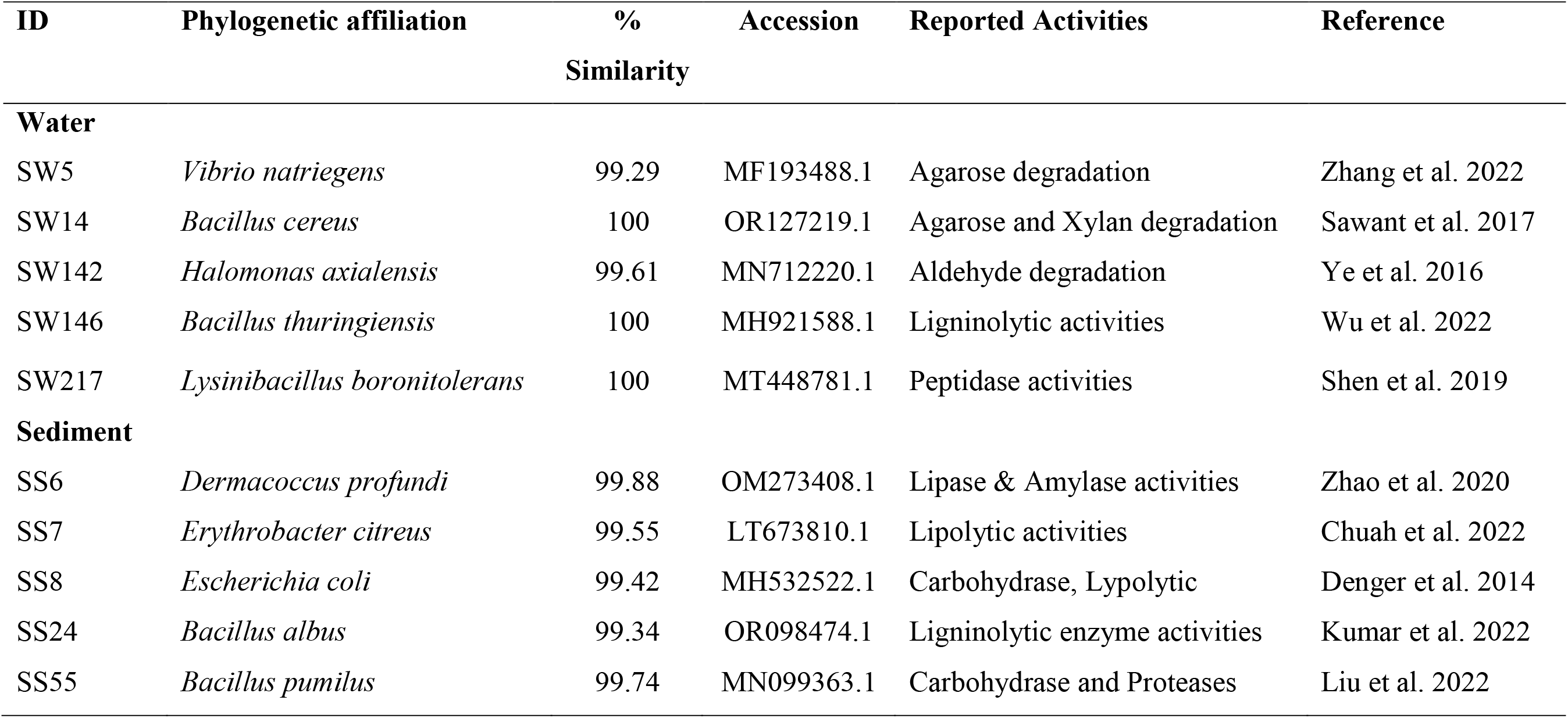
Bacterial morphotypes having the highest levels of enzyme activity from water and sediment, as well as their previously known enzyme activities in other marine environments.

